# Development of copy number assays for detection and surveillance of piperaquine resistance associated *plasmepsin 2/3* copy number variation in *Plasmodium falciparum*

**DOI:** 10.1101/655209

**Authors:** Christopher G Jacob, Megan R Ansbro, Roberto Amato, Mihir Kekre, Ranitha Vongpromek, Mehul Dhorda, Chanaki Amaratunga, Sokunthea Sreng, Seila Suon, Olivo Miotto, Rick M Fairhurst, Thomas E Wellems, Dominic P Kwiatkowski

## Abstract

Long regarded as an epicenter of drug-resistant malaria, Southeast Asia continues to provide new challenges to the control of *Plasmodium falciparum* malaria. Recently, resistance to the artemisinin combination therapy partner drug piperaquine has been observed in multiple locations across Southeast Asia. Genetic studies have identified a single nucleotide polymorphism as well as a copy number variation molecular marker that associate with clinical and *in vitro* resistance. The copy number polymorphism is a duplication of a region containing members of the plasmepsin multi-gene family of proteases. To accurately and quickly determine the presence of copy number variation in the *plasmepsin 2/3* duplication in field isolates, we developed a quantitative PCR assay using TaqMan probes. We validated copy number estimates using a separate SYBR green-based quantitative PCR assay as well as a novel breakpoint assay to detect the hybrid gene product. Field samples from 2012 – 2015 across 3 sites in Cambodia were tested using DNA extracted from dried blood spots and whole blood to monitor the extent of *plasmepsin 2/3* gene duplications, as well as *pfmdr1*. We found high concordance across all methods of copy number detection. For samples derived from dried blood spots we found a greater than 80% success rate in each assay, with more recent samples performing better. We found evidence of extensive plasmepsin 2/3 copy number amplifications in Pursat (94%, 2015) and Preah Vihear (87%, 2014), and lower levels in Ratanakiri (16%, 2014) in eastern Cambodia. We also see evidence of a shift from two copies of *plasmepsin 2/3* in Pursat 2013 to three copies in 2014-15 (25% to 64%). *Pfmdr1* duplications are absent from all samples in 2014 from Preah Vihear and Ratanakiri and 2015 from Pursat. This study shows increasing levels of *plasmepsin 2/3* gene amplifications across Cambodia from 2012 – 2015 and a complete reversion of *pfmdr1* mutant parasites in all study locations. The multiplex TaqMan assay is a robust tool for monitoring both *plasmepsin* and *pfmdr1* copy number variations in field isolates, and the SYBR-green and breakpoint assays are useful for monitoring *plasmepsin 2/3* duplications.

## Background

As malaria endemic countries strive toward malaria elimination, one of the main obstacles is the continued availability of efficacious drugs. In Southeast Asia, drug resistance is wide-spread with the most recent emergence to artemisinin combination therapies (ACTs), which are currently used as first-line treatment regimens.[1] The declining efficacy of ACTs in recent years can be largely attributed to rising resistance to artemisinin partner drugs, notably piperaquine. Dihydroartemisinin-piperaquine (DHA-PPQ) was introduced as the first-line treatment for malaria in 2008 in Cambodia, following partner drug resistance to artesunate-mefloquine (AS-MQ), the ACT used prior to 2008. Since 2012-2013, studies in Cambodia have shown declining efficacy to piperaquine *in vitro*, and subsequent increases in clinical treatment failures [2-7]. Genomic studies carried out in parallel with samples from these clinical efficacy studies have shown that there are multiple signals across the parasite genome that associate with both *in vitro* piperaquine resistance and clinical treatment failures[8, 9]. Specifically, a gene duplication within the *plasmepsin* multi-gene cluster on the parasite chromosome 14 and a non-synonymous SNP in a putative exonuclease gene (PF3D7_1362500) on chromosome 13, *exo-E415G*. Additional work also points to mutations in the chloroquine resistance transporter gene, *pfcrt* that can confer differing levels of piperaquine resistance in field and lab isolates [10, 11].

The duplication encompasses *plasmepsin 2* and a hybrid of the *plasmepsin 1 and 3* genes and was highly correlated (adjusted hazard-ratio 16.7) with parasite recrudescence following adequate drug treatment with DHA-PPQ. This effect holds in the artemisinin resistance associated *kelch13* propeller <u>(K13)</u> domain mutant populations [12] (adjusted hazard-ratio 5.2) suggesting an independent mechanism [8] Members of the *plasmepsin* gene family in *P. falciparum* are involved in the hemoglobin degradation pathway, specifically in the formation of hemozoin[13]. As the parasites digest hemoglobin and release heme, toxic by-products that cause oxidative stress are formed and the conversion of intermediates to inert hemozoin crystals detoxifies the harmful byproducts in the hemoglobin digestion pathway. The plasmepsin enzymes are redundant and other enzymes also facilitate the hemoglobin digestion pathway, including facilysins and falcipains. [14-16]

The main duplication in *plasmepsin 2/3* observed in Southeast Asia has a conserved break-point within the distal end of *plasmepsin 3* and includes complete duplication of the *plasmepsin 2* gene. In the same studies, an association was seen between increased copy numbers of *pfmdr1* and low piperaquine IC_50_, and very few cases where both *plasmepsin 2/3* and *pfmdr1* were duplicated within the same parasite. It is unknown if the association between decreased piperaquine IC_50_ and single copy *pfmdr1* is due to a drug effect, or is due to the expansion of piperaquine resistance on a mefloquine sensitive parasite line. With the observation of possible counteracting resistance mechanisms, it has been suggested to re-introduce mefloquine to areas of emerging piperaquine resistance, or to combine mefloquine into a triple ACT (TACT) with piperaquine. Both options are currently being investigated[17].

Monitoring this molecular marker of piperaquine resistance is essential to determine the frequency in populations which use DHA-PPQ as well as to determine if it appears in new parasite populations. We developed a TaqMan based quantitative PCR (qPCR) to measure the copy number of *plasmepsin 2* within the duplicated region, and have also combined it with a TaqMan assay that can detect increased copy numbers of *pfmdr1*. This single reaction multiplex qPCR can be used to efficiently monitor resistance to both piperaquine and mefloquine, a desirable prospect as mefloquine is in discussion for re-introduction to Cambodia in TACTs. We have also developed a PCR based breakpoint assay for detection of the hybrid sequence created as a result of the *plasmepsin 2/3* duplication. This assay can be used in in conjunction with the qPCR assays and/or in low-resource settings where qPCR is infeasible.

## Methods

### Samples

Laboratory isolates used in qPCR validation were obtained from a clinical trial carried out in 3 sites in Cambodia between 2012 and 2013[2]. Blood samples from this study were taken as whole-blood venous draws following malaria diagnosis and from an initial finger prick dried blood spot (DBS). A subset of DNA samples extracted from the venous blood were whole-genome sequenced and *pfmdr1* and *plasmepsin 2* copy-numbers were called from sequence data according to Amato et al.[8]. Additional field derived isolates (after 2013) from clinical trials performed by the NIH at the same 3 sites in Cambodia as above were used to test ongoing copy-number polymorphisms (clinicaltrials.gov ID: NCT01736319).

### Quantification of *plasmepsin 2, plasmepsin3*, and *pfmdr1* by real-time PCR

Primers for both *plasmepsin 2* and *plasmepsin 3* were designed using GenScript online tool (https://www.genscript.com/tools/) to match the T_m_ of the previously described *pfmdr1* TaqMan assay[18]. *Plasmepsin 2* primers (forward *-* 5’– ATGGTGATGCAGAAGTTGGA-3’, reverse - 5’–AACATCCTGCAGTTGTACATTTAAC-3’) and *plasmepsin 3* primers (forward - 5’–CCACTTGTGGTAACACGAAATTA-3’; reverse - 5’–TGGTTCAAGGTATTGTTTAGGTTC-3’) were selected to match the optimized reaction conditions of the *pfmdr1* primers (forward 5’-TGCATCTATAAAACGATCAGACAAA-3’, reverse 5’-TCGTGTGTTCCATGTGACTGT-3’. For the *plasmepsin* assay the same β*-tubulin* single copy reference primers (forward 5’ –TGATGTGCGCAAGTGATCC-3’, reverse 5’–TCCTTTGTGGACATTCTTCCTC-3’) were used as in the *pfmdr1* assay. First, the two-probe assay with either *plasmepsin 2* (5’Fam–CAGGATCTGCTAATTTATGGGTCCCA-3’BHQ-2) or *plasmepsin 3* (5’Fam– CCAACACTCGAATATCGTTCACCAA-3’BHQ-2) and beta-tubulin (5’MAX– TAGCACATGCCGTTAAATATCTTCCATGTCT-3’BHQ-1) was validated first using DNA from whole-blood extracted DNA and compared with copy-number estimates called from whole-genome sequence data. The *plasmepsin 2* probe set was then multiplexed with *pfmdr1* (5’ Cy5-TTTAATAACCCTGATCGAAATGGAACCTTTG-3’BHQ-2) and then tested using DNA extracted from DBS in the same set of samples. All primers and probes were ordered from Integrated DNA Technologies, Inc.. Reactions were carried out in 25μl volumes in 96 well plates (Star Labs) on a Roche 480 LightCycler. For each reaction we used 2X Concentrated Roche LightCycler 480 Probes Master (containing FastStart Taq DNA Polymerase), 300 nmol/L of each *plasmepsin 2* or *plasmepsin 3, pfmdr1*, and β*-tubulin* primers, 100 nmol/L of each probe, and 2-5 μl of sample DNA. PCR cycling conditions had an initial step of 95 °C for 10 minutes followed by 50 cycles of 95°C for 15 seconds and 58°C for 60 seconds. To calculate the fold-change we used the Δ ΔCt method, Δ ΔCt = (Ct_TE_ – Ct_HE_) – (Ct_TC_ – Ct_HC_) where T is the test gene (either *plasmepsin 2, plasmepsin 3*, or *pfmdr1*), H is the reference gene (β*-tubulin*), E is the experimental sample, and C is the control sample (for all tests we used 3D7 as the single copy control). We calculated relative expression as 2^-ΔΔCt.^

### SYBR green validation of *plasmepsin 2* copy number using quantitative PCR

PCR primers for estimating *plasmepsin 2/3* copy number amplification were designed manually inside the *plasmepsin 2* gene (forward - 5’-CTTATACTGCTTCAACATTTGATGGTATCCTTGG-3’; reverse - 5’-GGTTAAGAATCCTGTATGTTTATCATGTACAGGTAAG-3’). Previously described primers for *P. falciparum lactate dehydrogenase* (*ldh*)[19], were used as the control for a single copy gene (forward - 5’-AGGACAATATGGACACTCCGAT-3’; reverse - 5’-TTTCAGCTATGGCTTCATCAAA-3’). Quantitative PCR reactions were carried out in 20 μl volumes in a 96-well plate (Bio-Rad, Hercules, CA) containing 10 μl SensiFAST SYBR No-ROX mix (2x) (Bioline Inc.,Taunton, MA), 300 nM of each primer, and 2 μl genomic DNA. Reactions were performed using a CFX Connect Real-Time PCR Detection System (Bio-Rad) using the following conditions: 5 minutes at 95°C, followed by 40 cycles of 10 seconds at 95°C, 20 seconds at 58°C, and 20 seconds at 60°C. Relative copy number was calculated on the basis of the 2^-ΔΔCt^ method for relative quantification. ΔΔCt was calculated as (Ct_*ldh*_ –Ct_*pfplasmepsin2*_) - (Ct_*ldh* cal_ -*Ct*_*pfplasmepsin2* cal_), where cal is the calibration control of genomic 3D7 DNA with one copy of both *ldh* and *plasmepsin2*. DNA from an isolate with two copies of *plasmepsin 2/3* (PH1387-C)[2] was used as an internal plate control. All samples were analyzed in triplicate and each plate was replicated in triplicate.

### Plasmepsin 2/3 Duplication Breakpoint PCR Assay

Whole genome sequencing (WGS) data from *P. falciparum* genomic DNA collected during field studies in Cambodia was used to detect the *plasmepsin 2/3* gene amplification as previously reported[2, 8]. With the available WGS data, the breakpoint of the *plasmepsin 2/3* amplification was used to manually design PCR primers to amplify the region surrounding the breakpoint. Primers AF (forward 5’-CCACGATTTATATTGGCAAGTTGATTTAG-3’) and AR (reverse 5’-CATTTCTACTAAAATAGCTTTAGCATCATTCACG-3’) amplify a 623 bp product surrounding the breakpoint located at the 3’ end of *plasmepsin 1*. Primers BF (forward - 5’-CGTAGAATCTGCAAGTGT TTTCAAAG-3’) and BR (reverse 5’-AATGTTATAAATGCAATATAATCAAACGACATCAC-3’) amplify a 484 bp product surrounding the breakpoint located at the 3’ end of *plasmepsin 3*. BF + AR amplify the junction between the breakpoint and produce a 497 bp product in isolates with *plasmepsin 2/3* amplifications. These primers face opposite directions in samples without duplications and are not expected to amplify a product in single copy isolates. Both control (AF + AR; BF+ BR) and duplication (BF + AR) primer sets were used with all samples and one copy isolates were only noted if the control primer sets amplified a product and duplication PCR was negative. Two or more copies were annotated as >1 copy of *plasmepsin 2/3* only if both the control and duplication primer sets produced a product. PCR reactions contained 10 μl SapphireAmp Fast PCR Master Mix (Takara Bio USA, Mountain View, CA), 0.3 μl of each primer (10 μM stocks), 1 μl of genomic DNA up to 20 μl final volume with water. PCR conditions were: 92°C for 2 minutes, followed by 30 cycles of 92°C for 30 seconds, 59°C for 30 seconds, 66°C for 1.5 minutes, followed by a 1 minute extension at 66°C.

## Results

### Validation of copy number assays

Assays for *plasmepsin 2* and *plasmepsin 3* were designed and compared to determine if both genes could serve as markers for the entire duplication. For surveillance purposes the *plasmepsin 2* assay was multiplexed with an existing *pfmdr1* TaqMan assay. Both reference laboratory strains and culture-adapted field isolates were used to test the repeatability of each assay. The 3D7 isolate has single copies of all genes being tested while the Dd2 parasite has a duplication of the *pfmdr1* gene and the field isolates PH1265-C and PH1387-C have duplications of the *plasmepsin 2/3* complex. Two field isolates (PH1097-C and PH1310-C) have single copies of all genes and were used as baseline controls. Individual *plasmepsin* assays and the combined *plasmepsin 2* – *pfmdr1* assay showed high replicability across duplicates (Table 1).

**Table 1.** Validation of TaqMan assays with laboratory isolates.

| Strain | Assay | Replicates | Plasmepsin Est. FC |  | MDR-1 Est. FC |  |
| --- | --- | --- | --- | --- | --- | --- |
|  |  |  | Avg | SD (range) | Avg | SD (range) |
| 3D7 | *plasmepsin 2* | 8 | 1.00 | 0.02 (0.97-1.04) | - | - |
| Dd2 | *plasmepsin 2* | 8 | 1.06 | 0.03 (1.03-1.12) | - | - |
| PH1097-C | *plasmepsin 2* | 6 | 1.06 | 0.02 (1.04-1.09) | - | - |
| PH1265-C | *plasmepsin 2* | 6 | 1.89 | 0.02 (1.85-1.93) | - | - |
| PH1310-C | *plasmepsin 2* | 6 | 1.06 | 0.03 (1.01-1.13) | - | - |
| PH1387-C | *plasmepsin 2* | 6 | 1.93 | 0.09 (1.8-2.04) | - | - |
| 3D7 | *plasmepsin 3* | 8 | 1.02 | 0.03 (0.96-1.08) | - | - |
| Dd2 | *plasmepsin 3* | 8 | 1.09 | 0.03 (1.05-1.15) | - | - |
| PH1097-C | *plasmepsin 3* | 6 | 1.06 | 0.02 (1.03-1.09) | - | - |
| PH1265-C | *plasmepsin 3* | 6 | 2.05 | 0.06 (1.93-2.14) | - | - |
| PH1310-C | *plasmepsin 3* | 6 | 1.00 | 0.03 (1.02-1.09) | - | - |
| PH1387-C | *plasmepsin 3* | 6 | 2.29 | 0.07 (2.19-2.38) | - | - |
| 3D7 | *plasmepsin 2 / pfmdr1* | 8 | 1.01 | 0.02 (0.97-1.04) | 1.00 | 0.02 (0.96-1.02) |
| Dd2 | *plasmepsin 2 / pfmdr1* | 8 | 1.10 | 0.04 (1.04-1.17) | 1.83 | 0.06 (1.77-1.92) |
| PH1097-C | *plasmepsin 2 / pfmdr1* | 6 | 1.07 | 0.05 (1-1.13) | 1.06 | 0.05 (1.01-1.11) |
| PH1265-C | *plasmepsin 2 / pfmdr1* | 6 | 1.80 | 0.03 (1.75-1.84) | 0.96 | 0.02 (0.93-0.99) |
| PH1310-C | *plasmepsin 2 / pfmdr1* | 6 | 1.02 | 0.02 (1-1.06) | 1.01 | 0.02 (0.98-1.05) |
| PH1387-C | *plasmepsin 2 / pfmdr1* | 6 | 1.89 | 0.05 (1.84-2) | 0.93 | 0.03 (0.91-0.99) |

We then tested each assay from DNA extracted from whole blood for 67 patient samples. Copy number estimates from both the *plasmepsin 2* and *plasmepsin 3* assays were compared and were in 100% concordance across all samples extracted from venous blood. Fold-change values were comparable across both assays. Fifty-six of the 67 tested samples had WGS data available and had *plasmepsin 2/3* copy-number estimates available; 53 of the 56 samples with copy-number estimates from WGS had the same estimate from qPCR, with the discrepancies being 1 sample called 2 copies by qPCR and 3 copies by WGS and 2 samples with 4 copies by qPCR and 3 copies by WGS. WGS estimates were used to define the relative expression boundaries between copy number estimates, and relative-expression values were lower in whole-blood extracted samples than in cultured laboratory isolates (Figure 1A). We observed a high proportion of the *exo-E415G* mutation among *plasmepsin 2* multi-copy parasites (Figure 1). *Exo-415* calls were extracted from samples with available WGS data[8].

**FIG 1.**
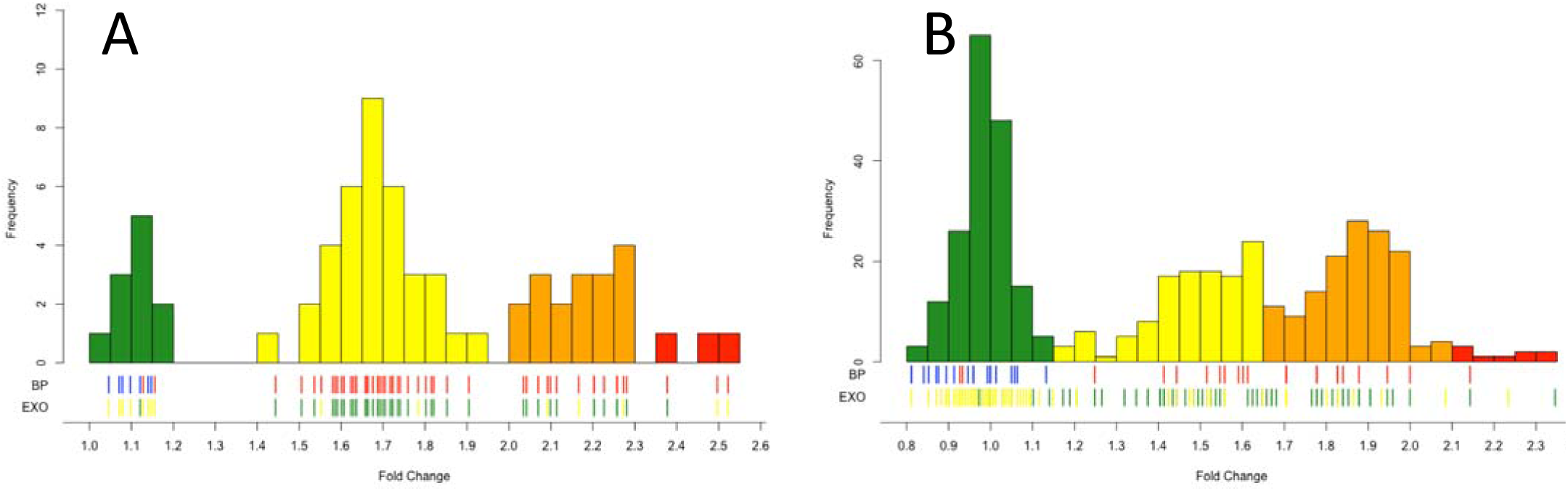
Distribution of fold-changes of samples from whole-blood (A) or dried blood spots (B). Tick marks indicate individual samples status for breakpoint (BP), with blue being no breakpoint detected and red having the breakpoint, or exo marker, where yellow is and E allele and green is the G allele. Bars are colored by their predicted *plasmepsin2/3* copy number of either 1, 2, 3, or 4+ (green, yellow, orange, and red respectively).

To validate the copy number estimates, available samples were tested at the National Institutes of Health (NIH) in the Laboratory of Malaria and Vector Research (LMVR) in Bethesda, MD, USA using a separately developed SYBR-green based approach. Of the 31 samples available at both laboratories, 29 samples matched in copy number estimate with one sample being called 2 copies by TaqMan and 3 copies by SYBR-green, and one sample being called 3 copies by TaqMan and 2 copies by SYBR-green. All dual-tested samples that had copy number estimates from both the SYBR-green method and WGS were in 100% concordance.

### Duplication breakpoint assay validation

We designed PCR primers that amplify the breakpoint of the *plasmepsin 2/3* amplification observed in Cambodian isolates (Figure 2A). Our breakpoint assay identifies copy number amplification in isolates that contain 2 or greater copies of *plasmepsin 2* and *3* (Figure 2B). As expected, no PCR products were observed in samples with a single copy of *plasmepsin 2/3* (Figure 2B). Control primers confirmed that the regions surrounding the *plasmepsin 2/3* amplification were present in all isolates (Figure 2C).

**FIG 2.**
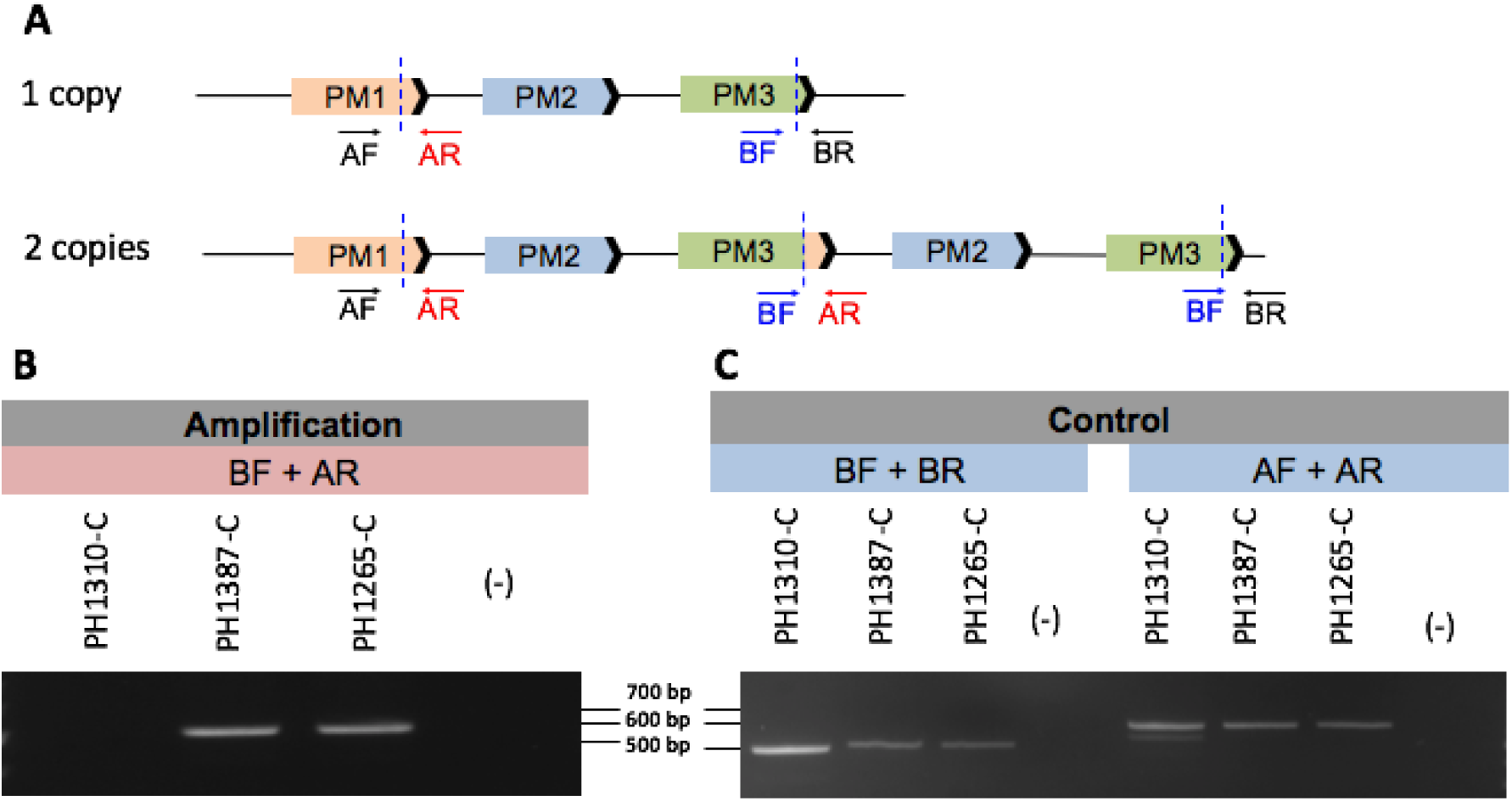
Schematic of *plasmepsin 2/3 gene* duplication. A) Gene model depicting the *plasmepsin 2/3* breakpoint (dashed blue lines) observed in Cambodian isolates. Primer positions are labeled in the single copy (top) and multi-copy (bottom) isolates. B. Amplification primer set BF + AR amplifies a product in an isolate with two copies (PH1387-C) and three copies (PH1265-C) of *plasmepsin 2/3*. No product is observed for the single copy (PH1310-C) isolate or in the DNA-negative control (-). C. Control primers amplify a product in the single copy (PH1310-C) and multi-copy isolates (PH1387-C; PH1265-C). No product is observed in the DNA-negative control (-).

Further verification of the breakpoint assay was performed via sequence analysis of the BF + AR PCR product. Sequence data revealed the same breakpoint location as observed by WGS [8]. All Cambodian samples that were positive for *plasmepsin 2/3* amplification as detected by the breakpoint assay (>1 copy) were in 100% concordance with qPCR and WGS data that called 2 or more copies of *plasmepsin 2*. Sequencing chromatogram review of PCRs representing 2 and 3 copy samples showed that the breakpoints for representative 2 and 3 copy samples were identical. These sequencing results combined with the identical PCR sizes for all isolates indicates that the breakpoint is identical in all Cambodian isolates tested to date.

To test the utility of using the breakpoint assay for rapid surveillance of large sample sizes for which WGS data is not yet completed or available, we analyzed the presence of amplification in an additional 99 samples. We found that 93/99 (94%) samples tested with the breakpoint PCR assay matched qPCR data for the same samples. The six non-concordant samples were repeated, always producing the same results with the breakpoint detecting a copy-number increase and the qPCR single-copy. These results suggest that the breakpoint PCR assay is more sensitive than the qPCR assay for detecting minor clones containing the duplication in field isolates.

### Copy number surveillance in Cambodia

To test the feasibility of using this assay as a surveillance tool, we performed the *plasmepsin 2* - *pfmdr1* TaqMan assay on 524 samples extracted from DBS collected in drug efficacy studies from 3 field sites within Cambodia (Pursat, Preah Vihear, & Ratanakiri) from 2012 to 2015. There was an 84% (success rate across all samples, but success rate improved with time since collection (2012, 68%; 2013, 72%; 2014, 94%; 2015, 100%). Assays run on samples from DBS were checked against SYBR-green results run on whole-blood extracted samples analyzed at the NIH LMVR. A total of 171 samples were available for analysis using both qPCR assays and had an 88% overall concordance, with only 6 (3%) samples having a discrepancy between calling a single versus multi-copy parasite. Most differences were between calling 2 versus 3 copies in either assay. Additionally, 71 samples that were either unavailable for DBS extraction or failed TaqMan qPCR were able to be typed by the SYBR-green method, giving copy-number estimates for 509 total samples. Samples with qPCR estimates in multiple sample types (whole-blood and DBS) were used to define relative expression limits between number of copies. We noticed a drop in relative expression in samples from DBS compared to whole-blood samples but confirmed multiple copies by the break-point assay, as well as observing *exo-E415G* SNPs among multi-copy samples (Figure 1B).

We observed an increase in multi-copy *plasmepsin 2* each year in both Pursat and Preah Vihear, but did not observe any multi-copy containing parasites in Ratanakiri until 2014 when they were observed in 17% of parasites analyzed. We also observed an increase in parasites containing 3 copies of *plasmepsin 2* from 2012/13 to 2014/15 to where they are now the majority (64%). There was also a large increase in multi-copy containing parasites in Preah Vihear from 2013 (29%) to 2014 (87%) (Figure 3A). Among all 438 samples with *pfmdr1* copy-number estimates, only 10 (2%) had multiple copies and no parasites with duplications were seen in the most recently sampled year for each site (Figure 3B).

**FIG 3.**
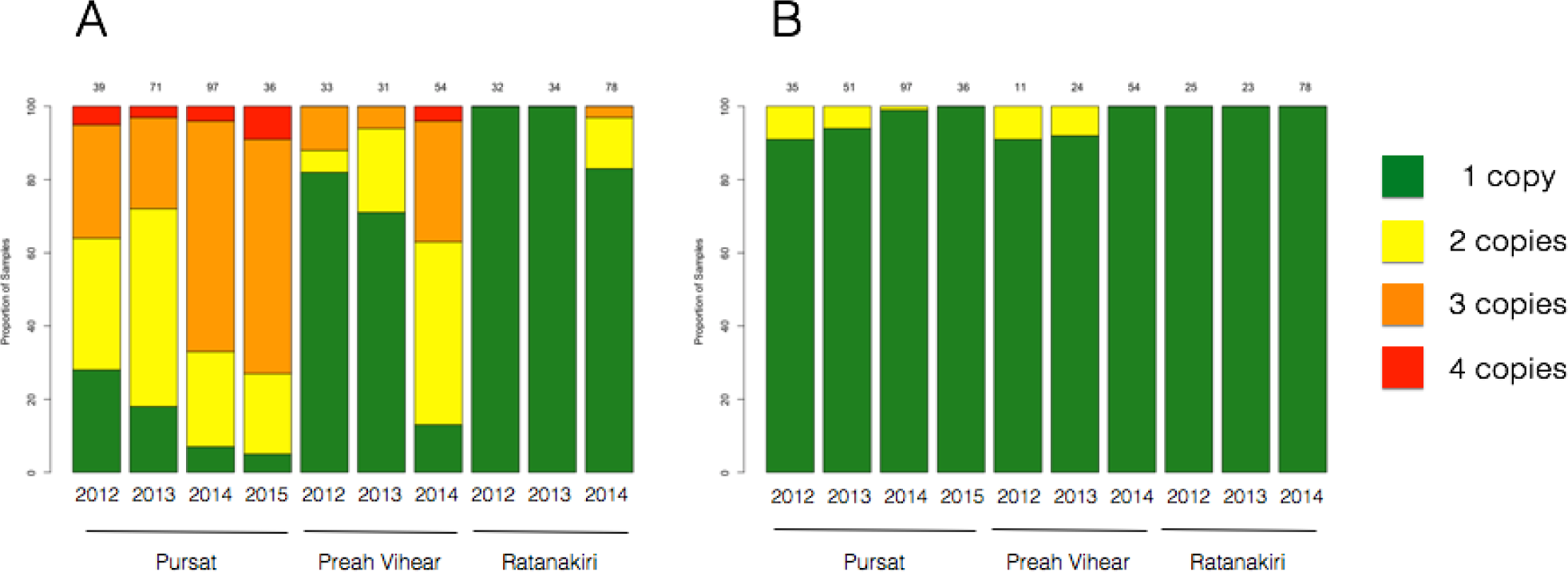
Proportion of samples by copy-number of *plasmepsin* (A) and *mdr1* (B). Bars represent proportion of samples by site and year for assays of dried-blood spot derived samples, and are colored by number of copies detected.

## Discussion

Our newly developed TaqMan assay accurately determines *plasmepsin 2/3* and *mdr1* gene amplifications in samples from DBS and can be performed using standard lab equipment and a suitable qPCR machine. The specificity of TaqMan probes combined with the different absorbance spectra of labeling dyes create a system for typing the reference gene simultaneously with one or more experimental genes, making it higher throughput than SYBR-green based methods. Our method shows high typeability in samples collected on dried filter paper blood spots making it ideal for surveillance of populations across wide areas. The breakpoint PCR assay also effectively determines increased *plasmepsin 2/3* copy number in field isolates and can be used in areas where qPCR is infeasible. The sensitivity of this assay enables the detection of minor clones, which proves advantageous, given the potential for polyclonal infections. However, a notable disadvantage of this method is the inability to distinguish the relative copy number (2, 3, or 4+ copies vs. >1) in comparison with qPCR. The breakpoint assay was also designed with primers specific to the *plasmepsin 2/3* breakpoint observed in Cambodian field isolates[8]. The PCR primers used to detect the Cambodian breakpoint observed to date may not amplify a product in samples that contain different breakpoints. As previous studies have suggested[20, 21], copy number variations in the same gene can arise independently on different genetic backgrounds and result in distinct molecular breakpoints of amplification. Since qPCR methods do not rely on the location of primers in reference to an estimated breakpoint, they can be used broadly. Furthermore, the high-throughput potential of the qPCR assay to type multiple genes in the same assay while providing actual copy number estimates makes the TaqMan assay the preferred method in areas where qPCR is possible.

Since the first *Plasmodium falciparum* genome was sequenced in 2002 [22], nearly 10,000 additional genomes have been sequenced from parasites collected around the endemic world. While the first genome may not have made great strides to discover new drug or vaccine targets or point towards complicated mechanisms of disease, it was the first step in understanding the great complexity of the *Plasmodium* genomic landscape. It is this understanding that has led to great advances in determining new molecular markers of drug resistance. Historically, it has taken decades to determine a molecular cause of drug resistance as was the case with chloroquine[23]. The availability of new methods and a catalogue of genomic variation allowed for rapid discovery and publishing of a candidate molecular marker of artemisinin resistance in 2014 [12], shortly after reports of suspected drug resistance to artemisinin in Southeast Asia were published in 2008 [24] and 2009 [25].

In 2014 the first report of DHA-PPQ treatment failures was published[7]. Because of an ever-increasing catalogue of sequenced samples and clinical and laboratory studies, a putative marker of drug resistance was published less than two years after the first reports of resistance[8, 9], with additional studies identifying new markers in *pfcrt* shortly after [10, 11]. Being able to extensively type variation by WGS made it possible to identify the association of increased piperaquine IC_50_ values with both SNPs and a copy-number variation. The copy-number variation at the *plasmepsin 2/3* locus showed high correlation with the phenotype and the new assays for detection will assist in monitoring its frequency in established populations, and can also monitor unaffected populations to check for new emergence or spread.

For a molecular marker to be effective in surveillance it must be easily typed in field-derived samples. Most molecular markers of *P. falciparum* drug resistance are SNPs and can be typed by simple PCR assays[26-31], although copy-number markers exist for mefloquine[18] and antifolates[32] they are more difficult to type. Quantitative PCR makes for an easy and inexpensive method to determine copy number in samples, and unlike breakpoint assays, qPCR can determine the number of copies. It is not known if more than 2 copies of *plasmepsin 2/3* have a phenotypic effect but the increase in samples with 3 or more copies suggest some sort of selective mechanism. It is possible that a third copy of *plasmepsin 2/3* prevents a loss of resistance if the duplication is unstable and an extra copy is lost, therefore the third copy would act as a “buffer.

Recent studies in *Plasmodium* have provided insight into the functional role of the *plasmepsins* in response to piperaquine pressure. Loesbanluechai *et al*. 2018 showed that overexpression of *plasmepsin 2* and *plasmepsin 3* in the 3D7 parasite background did not change parasite susceptibility to piperaquine, artemisinin, or chloroquine[33]. Other studies have suggested that *plasmepsin 2* and *plasmepsin 3* knockouts in the same 3D7 background showed decreased piperaquine survival as measured by IC50 values[34]. Thus, additional functional work is needed to fully understand the relevance of the *plasmepsin 2/3* gene amplification in conferring any survival or fitness advantages in response to PPQ pressure.

Mefloquine is currently being considered for re-introduction as a partner drug for artemisinin either as a dual or triple combination[17]. It was removed as the first line treatment following widespread resistance via a duplication in the *pfmdr1* gene. Since mefloquine’s removal as a nationally-recommended treatment the levels of *pfmdr1* multi-copy number parasites has fallen and now is no longer detected in our samples from 3 distant sites in Cambodia (Figure 3). It has been suggested that a counter-acting mechanism of action has selected against multiple copies of *pfmdr1* in parasites subjected to piperaquine. This is feasible but will require confirmation, while another possibility is that the *plasmepsin 2/3* multi-copy parasites emerged on a *pfmdr1* single copy parasite background and it is this lineage(s) that have expanded.

In order to effectively monitor the spread of antimalarial drug resistance, it is imperative to have robust, high-throughput methods for detecting genetic markers of resistance. Accurate and timely surveillance of drug resistance markers informs treatment strategies and aids in maintaining and prolonging efficacy of the limited selection antimalarial drugs available. Our TaqMan qPCR and breakpoint assays are tools that can be widely used to track *plasmepsin 2/3* amplification as a marker of piperaquine resistance. As mefloquine is being reintroduced in multiple Southeast Asian countries, our multiplex TaqMan qPCR assay should prove useful in monitoring for genetic markers of both mefloquine and piperaquine resistance.

## Abbreviations

ACT: Artemisinin combination therapy,
DBS: dried blood spot,
DHA-PPQ: dihydroartemisinin-piperaquine,
IC_50_: 50% inhibitory concentration,
LMVR: Laboratory of Malaria and Vector Research,
NIH: National Institutes of Health,
qPCR: quantitative polymerase chain reaction,
WGS: whole-genome sequencing.

## Declarations

### Ethics approval and consent to participate

All participants or guardians provided written consent and samples were collected under approval from the Cambodian National Ethics Committee for Health Research and the National Institute of Allergy and Infectious Diseases Institutional Review board.

### Availability of data and material

Data sharing is not applicable to this article as no datasets were generated or analyzed during the current study.

### Competing interests

The authors declare that they have no competing interests.

### Funding

This work was supported by the Wellcome Trust (090770/Z/09/Z; 098051), the Medical Research Council UK and the Department for International Development (DFID) (G0600718 and MR/M005212/1), the Intramural Research Program of the NIAID, and the Bill & Melinda Gates Foundation (OPP1118166).

### Authors’ contributions

CGJ designed TaqMan assays and analyzed data. MRA designed SYBR-green and breakpoint assays and analyzed data. RA and OM provided CNV estimates from WGS data. MK, RV, & MD processed DBS samples. CA, SSr, SSu, & RMF conducted the clinical trials and collected the samples for this study. CGJ and MRA wrote the manuscript and prepared figures. RMF, TEW, and DPK edited figures and manuscript. All authors read and approved the final manuscript.

## Acknowledgements

The authors wish to thank the members of the Lee lab at the Sanger Institute for contributing DNA from laboratory isolates for experimental controls.

## References

1. Organization WGWH: World Malaria Report 2018. 2018.

2. Amaratunga C, Lim P, Suon S, Sreng S, Mao S, Sopha C, Sam B, Dek D, Try V, Amato R, et al: Dihydroartemisinin-piperaquine resistance in Plasmodium falciparum malaria in Cambodia: a multisite prospective cohort study. Lancet Infect Dis 2016, 16:357–365.

3. Chaorattanakawee S, Saunders DL, Sea D, Chanarat N, Yingyuen K, Sundrakes S, Saingam P, Buathong N, Sriwichai S, Chann S, et al: Ex Vivo Drug Susceptibility Testing and Molecular Profiling of Clinical Plasmodium falciparum Isolates from Cambodia from 2008 to 2013 Suggest Emerging Piperaquine Resistance. Antimicrob Agents Chemother 2015, 59:4631–4643.

4. Spring MD, Lin JT, Manning JE, Vanachayangkul P, Somethy S, Bun R, Se Y, Chann S, Ittiverakul M, Sia-ngam P, et al: Dihydroartemisinin-piperaquine failure associated with a triple mutant including kelch13 C580Y in Cambodia: an observational cohort study. Lancet Infect Dis 2015, 15:683–691.

5. Leang R, Barrette A, Bouth DM, Menard D, Abdur R, Duong S, Ringwald P: Efficacy of dihydroartemisinin-piperaquine for treatment of uncomplicated Plasmodium falciparum and Plasmodium vivax in Cambodia, 2008 to 2010. Antimicrob Agents Chemother 2013, 57:818–826.

6. Leang R, Taylor WR, Bouth DM, Song L, Tarning J, Char MC, Kim S, Witkowski B, Duru V, Domergue A, et al: Evidence of Plasmodium falciparum Malaria Multidrug Resistance to Artemisinin and Piperaquine in Western Cambodia: Dihydroartemisinin-Piperaquine Open-Label Multicenter Clinical Assessment. Antimicrob Agents Chemother 2015, 59:4719–4726.

7. Saunders DL, Vanachayangkul P, Lon C, Program Usammr, National Center for Parasitology E, Malaria C, Royal Cambodian Armed F: Dihydroartemisinin-piperaquine failure in Cambodia. N Engl J Med 2014, 371:484–485.

8. Amato R, Lim P, Miotto O, Amaratunga C, Dek D, Pearson RD, Almagro-Garcia J, Neal AT, Sreng S, Suon S, et al: Genetic markers associated with dihydroartemisinin-piperaquine failure in Plasmodium falciparum malaria in Cambodia: a genotype-phenotype association study. Lancet Infect Dis 2016.

9. Witkowski B, Duru V, Khim N, Ross LS, Saintpierre B, Beghain J, Chy S, Kim S, Ke S, Kloeung N, et al: A surrogate marker of piperaquine-resistant Plasmodium falciparum malaria: a phenotype-genotype association study. Lancet Infect Dis 2016.

10. Ross LS, Dhingra SK, Mok S, Yeo T, Wicht KJ, Kumpornsin K, Takala-Harrison S, Witkowski B, Fairhurst RM, Ariey F, et al: Emerging Southeast Asian PfCRT mutations confer Plasmodium falciparum resistance to the first-line antimalarial piperaquine. Nat Commun 2018, 9:3314.

11. Agrawal S, Moser KA, Morton L, Cummings MP, Parihar A, Dwivedi A, Shetty AC, Drabek EF, Jacob CG, Henrich PP, et al: Association of a Novel Mutation in the Plasmodium falciparum Chloroquine Resistance Transporter With Decreased Piperaquine Sensitivity. J Infect Dis 2017, 216:468–476.

12. Ariey F, Witkowski B, Amaratunga C, Beghain J, Langlois AC, Khim N, Kim S, Duru V, Bouchier C, Ma L, et al: A molecular marker of artemisinin-resistant Plasmodium falciparum malaria. Nature 2014, 505:50–55.

13. Banerjee R, Liu J, Beatty W, Pelosof L, Klemba M, Goldberg DE: Four plasmepsins are active in the Plasmodium falciparum food vacuole, including a protease with an active-site histidine. Proc Natl Acad Sci U S A 2002, 99:990–995.

14. Goldberg DE: Hemoglobin degradation. Curr Top Microbiol Immunol 2005, 295:275–291.

15. Wunderlich J, Rohrbach P, Dalton JP: The malaria digestive vacuole. Front Biosci (Schol Ed) 2012, 4:1424–1448.

16. Moura PA, Dame JB, Fidock DA: Role of Plasmodium falciparum digestive vacuole plasmepsins in the specificity and antimalarial mode of action of cysteine and aspartic protease inhibitors. Antimicrob Agents Chemother 2009, 53:4968–4978.

17. Maxmen A: Back on TRAC: New trial launched in bid to outpace multidrug-resistant malaria. Nat Med 2016, 22:220–221.

18. Price RN, Uhlemann AC, Brockman A, McGready R, Ashley E, Phaipun L, Patel R, Laing K, Looareesuwan S, White NJ, et al: Mefloquine resistance in Plasmodium falciparum and increased pfmdr1 gene copy number. Lancet 2004, 364:438–447.

19. Lim P, Dek D, Try V, Eastman RT, Chy S, Sreng S, Suon S, Mao S, Sopha C, Sam B, et al: Ex vivo susceptibility of Plasmodium falciparum to antimalarial drugs in western, northern, and eastern Cambodia, 2011-2012: association with molecular markers. Antimicrob Agents Chemother 2013, 57:5277–5283.

20. Hostetler JB, Lo E, Kanjee U, Amaratunga C, Suon S, Sreng S, Mao S, Yewhalaw D, Mascarenhas A, Kwiatkowski DP, et al: Independent Origin and Global Distribution of Distinct Plasmodium vivax Duffy Binding Protein Gene Duplications. PLoS Negl Trop Dis 2016, 10:e0005091.

21. Menard D, Chan ER, Benedet C, Ratsimbasoa A, Kim S, Chim P, Do C, Witkowski B, Durand R, Thellier M, et al: Whole genome sequencing of field isolates reveals a common duplication of the Duffy binding protein gene in Malagasy Plasmodium vivax strains. PLoS Negl Trop Dis 2013, 7:e2489.

22. Gardner MJ, Hall N, Fung E, White O, Berriman M, Hyman RW, Carlton JM, Pain A, Nelson KE, Bowman S, et al: Genome sequence of the human malaria parasite Plasmodium falciparum. Nature 2002, 419:498–511.

23. Plowe CV: The evolution of drug-resistant malaria. Trans R Soc Trop Med Hyg 2009, 103 Suppl 1:S11–14.

24. Noedl H, Se Y, Schaecher K, Smith BL, Socheat D, Fukuda MM, Artemisinin Resistance in Cambodia 1 Study C: Evidence of artemisinin-resistant malaria in western Cambodia. N Engl J Med 2008, 359:2619–2620.

25. Dondorp AM, Nosten F, Yi P, Das D, Phyo AP, Tarning J, Lwin KM, Ariey F, Hanpithakpong W, Lee SJ, et al: Artemisinin resistance in Plasmodium falciparum malaria. N Engl J Med 2009, 361:455–467.

26. Djimde A, Doumbo OK, Cortese JF, Kayentao K, Doumbo S, Diourte Y, Coulibaly D, Dicko A, Su XZ, Nomura T, et al: A molecular marker for chloroquine-resistant falciparum malaria. N Engl J Med 2001, 344:257–263.

27. Plowe CV, Djimde A, Bouare M, Doumbo O, Wellems TE: Pyrimethamine and proguanil resistance-conferring mutations in Plasmodium falciparum dihydrofolate reductase: polymerase chain reaction methods for surveillance in Africa. Am J Trop Med Hyg 1995, 52:565–568.

28. Gyang FN, Peterson DS, Wellems TE: Plasmodium falciparum: rapid detection of dihydrofolate reductase mutations that confer resistance to cycloguanil and pyrimethamine. Exp Parasitol 1992, 74:470–472.

29. Wang P, Brooks DR, Sims PF, Hyde JE: A mutation-specific PCR system to detect sequence variation in the dihydropteroate synthetase gene of Plasmodium falciparum. Mol Biochem Parasitol 1995, 71:115–125.

30. Khan B, Omar S, Kanyara JN, Warren-Perry M, Nyalwidhe J, Peterson DS, Wellems T, Kaniaru S, Gitonga J, Mulaa FJ, Koech DK: Antifolate drug resistance and point mutations in Plasmodium falciparum in Kenya. Trans R Soc Trop Med Hyg 1997, 91:456–460.

31. Plowe CV, Cortese JF, Djimde A, Nwanyanwu OC, Watkins WM, Winstanley PA, Estrada-Franco JG, Mollinedo RE, Avila JC, Cespedes JL, et al: Mutations in Plasmodium falciparum dihydrofolate reductase and dihydropteroate synthase and epidemiologic patterns of pyrimethamine-sulfadoxine use and resistance. J Infect Dis 1997, 176:1590–1596.

32. Nair S, Miller B, Barends M, Jaidee A, Patel J, Mayxay M, Newton P, Nosten F, Ferdig MT, Anderson TJ: Adaptive copy number evolution in malaria parasites. PLoS Genet 2008, 4:e1000243.

33. Loesbanluechai D, Kotanan N, de Cozar C, Kochakarn T, Ansbro MR, Chotivanich K, White NJ, Wilairat P, Lee MCS, Gamo FJ, et al: Overexpression of plasmepsin II and plasmepsin III does not directly cause reduction in Plasmodium falciparum sensitivity to artesunate, chloroquine and piperaquine. Int J Parasitol Drugs Drug Resist 2019, 9:16–22.

34. Mukherjee A, Gagnon D, Wirth DF, Richard D: Inactivation of Plasmepsins 2 and 3 Sensitizes Plasmodium falciparum to the Antimalarial Drug Piperaquine. Antimicrob Agents Chemother 2018, 62.

